# The nitrogen phosphotransferase regulator PtsN (EIIA^Ntr^) regulates inorganic polyphosphate production in *Escherichia coli*

**DOI:** 10.1101/2021.10.15.464621

**Authors:** Marvin Q. Bowlin, Abagail R. Long, Joshua T. Huffines, Michael J. Gray

## Abstract

Inorganic polyphosphate (polyP) is synthesized by bacteria under stressful environmental conditions and acts by a variety of mechanisms to promote cell survival. While the kinase that synthesizes polyP (PPK, enocoded by the *ppk* gene) is well known, little is understood about how environmental stress signals lead to activation of this enzyme. Previous work has shown that the transcriptional regulators DksA, RpoN (*σ*^54^), and RpoE (*σ*^24^) positively regulate polyP production, but not *ppk* transcription, in *Escherichia coli*. In this work, we set out to examine the role of the alternative sigma factor RpoN and nitrogen starvation stress response pathways in controlling polyP synthesis in more detail. In the course of these experiments, we identified GlnG, GlrR, PhoP, PhoQ, RapZ, and GlmS as proteins that affect polyP production, and uncovered a central role for the nitrogen phosphotransferase regulator PtsN (EIIA^Ntr^) in a polyP regulatory pathway, acting upstream of DksA, downstream of RpoN, and apparently independently of RpoE. However, none of these regulators appears to act directly on PPK, and the mechanism(s) by which they modulate polyP production remain unclear. Unexpectedly, we also found that the pathways that regulate polyP production vary depending not only on the stress condition applied, but also on the composition of the media in which the cells were grown before exposure to polyP-inducing stress. These results constitute substantial progress towards deciphering the regulatory networks driving polyP production under stress, but highlight the remarkable complexity of this regulation and its connections to a broad range of stress-sensing pathways.

**IMPORTANCE:** Bacteria respond to changes in their environments with a complex regulatory network that controls the expression and activity of a wide range of effectors important for their survival. This stress response network is critical for the virulence of pathogenic bacteria and for the ability of all bacteria to grow in natural environments. Inorganic polyphosphate (polyP) is an evolutionarily ancient and almost universally conserved stress response effector that plays multiple roles in virulence, stress response, and survival in diverse organisms. This work provides new insights into the connections between well characterized nitrogen starvation and cell envelope stress response signaling pathways and the production of polyP in *Escherichia coli*.

## INTRODUCTION

Inorganic polyphosphate (polyP) is a biopolymer of up to hundreds of phosphate units that is produced by organisms of all domains of life (1–3). In bacteria, polyP has been known to be required for stress response and virulence for decades (4–6), and recent work has begun to decipher molecular mechanisms by which bacterial polyP directly disrupts phagocytic cell functions important in the host immune response to bacterial infections (7–11). These results have led to a growing interest in polyP metabolism and the recent identification of a range of chemicals that inhibit the bacterial polyP kinases (PPKs) responsible for polyP synthesis as promising potential anti-virulence drug candidates (7, 12–15).

Despite this, relatively little is known about how polyP production is regulated in bacteria. In the model organism *Escherichia coli*, polyP is undetectable during exponential growth in rich medium, but is synthesized rapidly upon exposure to a variety of stress conditions, including severe oxidative stress, heat shock, salt stress, and multiple types of starvation stresses (16–18). Early work identified a few regulators in *E. coli* that affected polyP synthesis under different conditions, but did not establish the mechanisms by which these acted (16, 19, 20). In *E. coli*, PPK and the polyP-degrading enzyme exopolyPase (PPX) are encoded in a bicistronic operon (21) whose transcription does not increase upon stress treatment (17, 22, 23). However, our previous results have shown that transcriptional regulators are required for robust polyP production in response to a nutrient limitation stress in which exponential-phase cultures of *E. coli* are shifted from rich medium into phosphate-limited glucose minimal medium (16). These include the RNA polymerase-binding protein DksA (24) and the stress-responsive alternative sigma factors RpoE and RpoN (22). These observations led us to hypothesize that these transcription factors regulate the expression of genes or proteins responsible for directly activating PPK activity under stress conditions. However, the transcriptional response to nutrient limitation is complex, and transcriptomics have not allowed us to identify any such directly-acting regulators to date (22).

The experiments described in this paper were aimed at deciphering the role of RpoN-dependent genes in polyP regulation. RpoN is the *E. coli σ*^54^-family sigma factor, and is notable for requiring additional ATPase proteins for activation of transcription at specific promoters (25, 26). These bacterial enhancer binding proteins (bEBPs) control specific and well-defined regulons in *E. coli* (27), and we hypothesized that by determining which bEBP(s) were necessary for polyP production we would be able to identify specific RpoN-dependent polyP regulators.

In the course of testing this hypothesis, we discovered that the bEBP GlnG, involved in the classical nitrogen starvation response in *E. coli* (28), is a positive regulator of polyP synthesis, but that this activity and the effect of RpoN on polyP synthesis are suppressed by the presence of glutamine in the rich medium before nutrient limitation, growth conditions under which polyP is not being produced. Further studies revealed that glutamine had a significant effect on which regulatory pathways were required for polyP synthesis after nutrient limitation, and that this was true not only of RpoN and GlnG, but also of RapZ, a regulator of the peptidoglycan precursor synthesis enzyme GlmS (29). The cell envelope stress responsive regulators GlrR (30–32) and PhoP (33) both also impacted polyP production, acting as negative regulators. By examining known nitrogen-responsive regulatory systems in *E. coli*, we identified a key role for the nitrogen phosphotransferase regulator PtsN (EIIA^Ntr^) (34) as an activator of polyP synthesis that acts downstream of RpoN, upstream of DksA, and apparently independently of RpoE.

The results of these studies are new insights into the network of regulators involved in modulating polyP production in *E. coli* and an increased appreciation that media composition can impact not only the extent of polyP synthesis but also the regulators involved in controlling that synthesis. However, all of the regulators we have identified so far appear to act indirectly, and we have yet to identify the genes, proteins, or metabolites directly responsible for activation of PPK under stress conditions.

## RESULTS

### The bEBPs GlnG and GlrR influence polyP production

As we have previously reported (22), Δ*rpoN* mutant *E. coli* had a significant defect in polyP synthesis upon nutrient limitation stress (Fig. 1A). Expression of *rpoN* from a plasmid rescued this phenotype, but did not increase polyP production in a wild-type strain (Fig. 1A). RpoN-dependent promoters require bEBPs (25, 26), so to narrow down the identity of the RpoN-dependent gene(s) involved in polyP synthesis, we measured polyP production in mutants lacking each of the 11 bEBPs present in *E. coli* MG1655 (27) (Figs. 1B,C). Mutants lacking *glnG* (also known as *ntrC*)(28) had a significant defect in polyP production and mutants lacking *glrR* produced significantly more polyP than the wild-type (Fig. 1B). These phenotypes canceled each other out, and a Δ*glrR ΔglnG* double mutant produced an amount of polyP indistinguishable from the wild-type. No other bEBP mutations affected the extent of polyP synthesis under these conditions (Fig. 1C).

**FIG 1.**
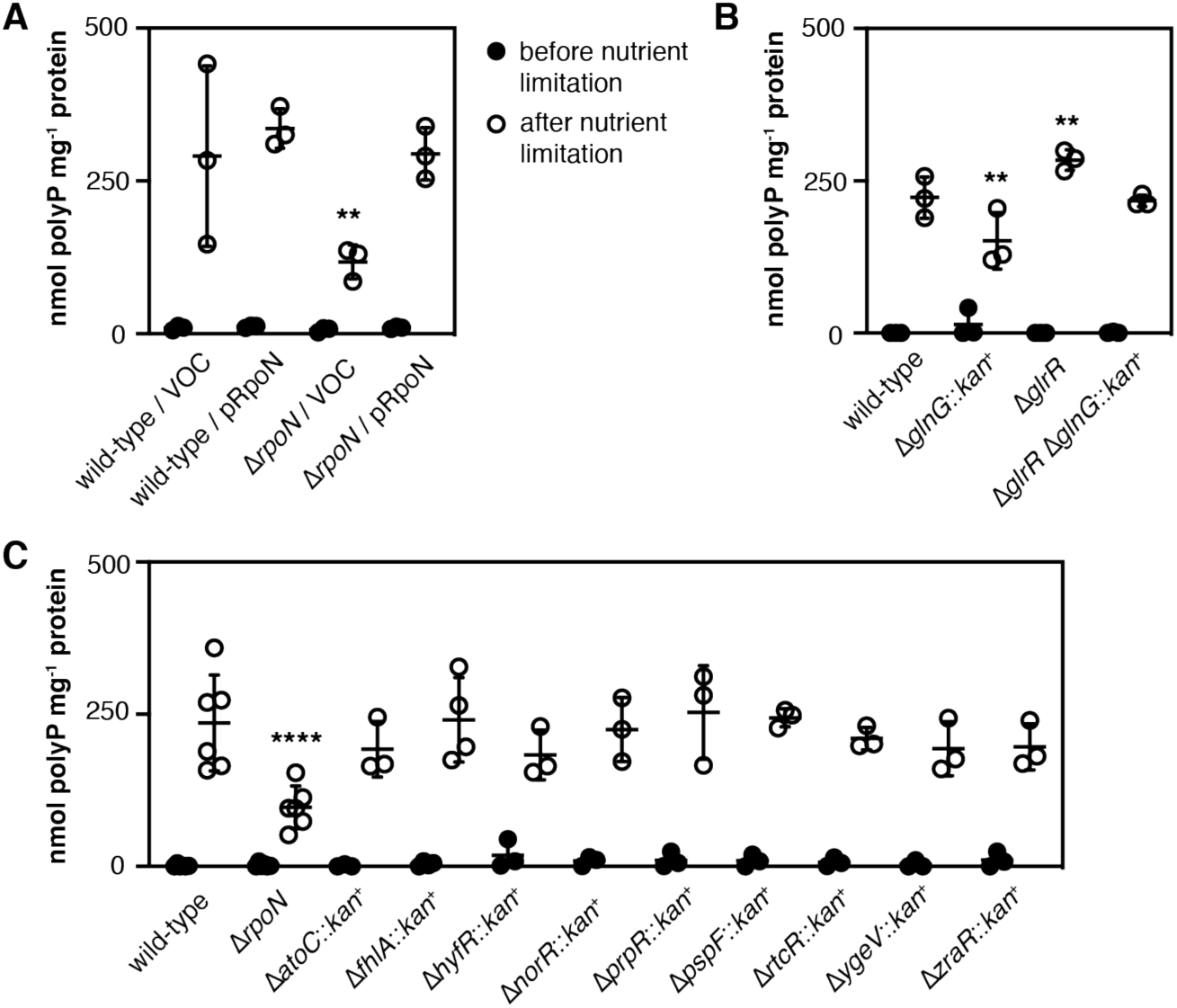
The bEBPs GlnG and GlrR influence polyP production. (*A*) *E. coli* MG1655 wild-type or Δ*rpoN730* containing either pBAD24 (VOC) or pRpoN (*rpoN*^+^) plasmids were grown at 37°C to A_600_=0.2–0.4 in LB containing 2 g l^-1^ arabinose (black circles) and then shifted to minimal medium containing 2 g l^-1^ arabinose for 2 hours (white circles)(n=3, ± SD). (*B*) MG1655 wild-type, Δ*glnG730*::*kan*^+^, Δ*glrR728*, or Δ*glrR728* Δ*glnG730*::*kan*^+^ were grown at 37°C to A_600_=0.2–0.4 in LB (black circles) and then shifted to minimal medium for 2 hours (white circles)(n=3, ± SD). (*C*) MG1655 wild-type, Δ*rpoN730*, Δ*atoC774*::*kan*^+^, Δ*hyfR739*::*kan*^+^, Δ*norR784*::*kan*^+^, Δ*prpR772*::*kan*^+^, Δ*pspF739*::*kan*^+^, Δ*rtcR755*::*kan*^+^, Δ*ygeV720*::*kan*^+^, or Δ*zraR775*::*kan*^+^ were grown at 37°C to A_600_=0.2–0.4 in LB (black circles) and then shifted to minimal medium for 2 hours (white circles)(n=3-6, ± SD). PolyP concentrations are in terms of individual phosphate monomers. Asterisks indicate polyP levels significantly different from those of the wild-type control for a given experiment (two-way repeated measures ANOVA with Holm-Sidak’s multiple comparisons test, ** = P<0.01, **** = P<0.0001).

### RpoN-dependent regulation of polyP synthesis is dependent on cellular nitrogen status, but not on GlnB or GlnK *per se*

The involvement of GlnG in polyP synthesis implicates the cellular response to nitrogen starvation in polyP regulation (28), which was not unexpected based on previous reports in the literature (16, 35). Under nitrogen limitation conditions, perceived by the cell as a decrease in the ratio of intracellular glutamine to glutamate and an accumulation of *α*-ketoglutarate (28, 36, 37), glutamine synthase (GlnA) is activated by a pathway involving the PII signaling proteins GlnB and GlnK (28, 38–41)(Fig. 2A). Transcription of *glnB* is constitutive, but *glnK* transcription is activated by RpoN and GlnG (28, 41, 42) (Fig. 2B), and *glnK* induction is a reliable reporter of cellular nitrogen starvation (41). Consistent with our previously-reported RNA sequencing data (22), polyP-inducing nutrient limitation strongly induced *glnK* expression (Fig. 2C). The gene encoding *glrR* is immediately adjacent to *glnB* in the *E. coli* genome, and the Δ*glrR* mutation used here deletes not only most of the coding sequence of that gene (43), but also deletes a binding site for the repressor PurR in the *glnB* promoter region (44, 45) (Fig. 2B). We therefore hypothesized that GlnK or GlnB, which regulate the activity of a variety of proteins by direct interaction (28, 38, 46–48), might be activators of polyP synthesis. To test this idea, we expressed *glnB* and *glnK* from arabinose-inducible plasmids, but found that neither *glnB* nor *glnK* overexpression increased polyP production in wild-type, Δ*rpoN*, or Δ*dksA* mutant strains (Figs. 2D,E). Mutants lacking both *glnB* and *glnK* are glutamine auxotrophs (49), and LB, the rich medium used for the “before stress” growth condition (16), is naturally very low in glutamine (50), so we tested whether Δ*glnB*, Δ*glnK*, or Δ*glnBK* mutations affected polyP production after nutrient shift from rich media supplemented with glutamine (LBQ; Fig. 2F). There was a very slight defect in polyP production in the Δ*glnB* mutant, but the more surprising result was that, although the extent of polyP production after shift from LBQ into minimal medium was very similar to that after shift from LB (Figs. 2D,E,F), neither *rpoN* nor *glnG* mutants had any defect in polyP synthesis under these conditions. This was unexpected and showed that cellular nitrogen status affects the regulatory pathway by which polyP synthesis is activated. This observation probably also explains the very high variability in polyP production we have previously reported in a Δ*glnG* mutant (24) in experiments performed with a different supply of LB medium. This was not true for the Δ*dksA* mutant (22, 24), which was defective in polyP synthesis regardless of whether the LB was supplemented with glutamine (Fig. 2F). Glutamine concentration is a common signal of cellular nitrogen status sensed by many regulators and enzymes (28, 37), so we tested whether PPK itself was allosterically regulated by either glutamate or glutamine, and found that neither of these compounds affected the activity of purified PPK *in vitro* at physiological concentrations (51) (Fig. 2G).

**FIG 2.**
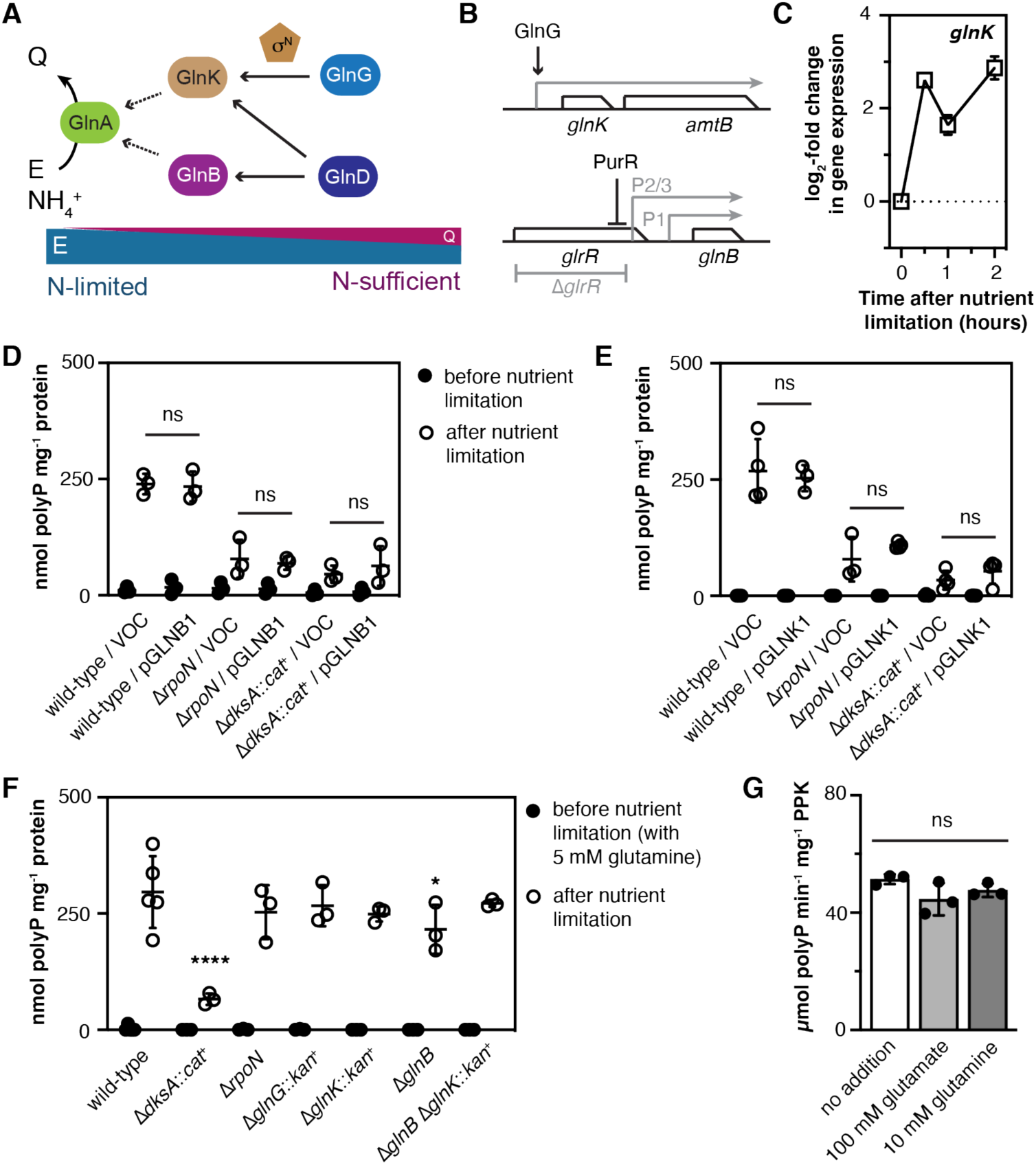
RpoN-dependent regulation of polyP synthesis is dependent on cellular nitrogen status, but not on GlnB or GlnK *per se*. (*A*) Simplified model of the regulation of GlnA (glutamine synthetase) activity in response to changes in the intracellular glutamate (E) to glutamine (Q) ratio. (*B*) Diagram of the *glnK* and *glnB* loci in *E. coli*. (*C*) *E. coli* MG1655 wild-type was grown at 37°C to A_600_=0.2–0.4 in rich medium (LB) and then shifted to minimal medium (MOPS with no amino acids, 4 g l^-1^ glucose, 0.1 mM K_2_HPO_4_, 0.1 mM uracil) for 2 hours. qRT-PCR was used to measure fold changes in *glnK* transcript abundance at the indicated timepoints (n=3, ± SD), normalized to expression before stress treatment (t = 0 h). (*D*, *E*) *E. coli* MG1655 wild-type, Δ*rpoN730*, or Δ*dksA1000*::*cat^+^* containing either pBAD18 (VOC), pGLNB1 (*glnB*^+^) or pGLNK1 (*glnK*^+^) plasmids were grown at 37°C to A_600_=0.2–0.4 in LB containing 2 g l^-1^ arabinose (black circles) and then shifted to minimal medium supplemented with 2 g l^-1^ arabinose for 2 hours (white circles)(n=3-4, ± SD). (*F*) MG1655 wild-type, Δ*dksA1000*::*cat^+^*, Δ*glnG730*::*kan*^+^, Δ*glnK736*::*kan*^+^, Δ*glnB727*, or Δ*glnB727* Δ*glnK736*::*kan*^+^ were grown at 37°C to A_600_=0.2–0.4 in LBQ (black circles) and then shifted to minimal medium for 2 hours (white circles)(n=3-5, ± SD). PolyP concentrations are in terms of individual phosphate monomers. Asterisks indicate polyP levels significantly different from those of the wild-type control for a given experiment (two-way repeated measures ANOVA with Holm-Sidak’s multiple comparisons test, ns = not significant * = P<0.05, ** = P<0.01, **** = P<0.0001). (*G*) *S*pecific activity of purified PPK in the presence of the indicated compounds (n=3, ±SD; one-way ANOVA, ns = not significant).

### Effect of *glmY* and GlmS (glutamine--fructose-6-phosphate aminotransferase) regulation on polyP production

Since we could not attribute the effect of the Δ*glrR* mutation on polyP synthesis (Fig. 1B) to disregulation of *glnB* (Fig. 2D), we examined the possible role of the GlrR regulon on polyP synthesis. GlrR is activated by GlrK, a histidine kinase that responds to cell envelope disruptions (30). In *E. coli* MG1655, GlrR is only known to regulate the expression of two promoters, that of the operon encoding the alternative sigma factor RpoE and that of the sRNA *glmY* (31, 32) (Fig. 3A). We have previously reported that Δ*rpoE* mutants have significant defects in polyP synthesis, indicating that RpoE is a positive regulator of polyP production (22). Regulation of the *rpoE* operon is complex (32, 52, 53), but GlrR is an activator of *rpoE* expression (32), so it is difficult to reconcile a model in which this explains the increase in polyP production in the Δ*glrR* mutant (Fig. 1B). We therefore turned our attention to *glmY*, which, in a pathway involving the RNA-binding protein RapZ and the sRNA *glmZ*, is responsible for increasing GlmS (glutamine--fructose-6-phosphate aminotransferase) synthesis under conditions where intracellular glucosamine-6-phosphate (GlcN6P) becomes limiting (29, 31, 54)(Fig. 3A). GlcN6P, synthesized by GlmS from glutamine and fructose-6-phosphate, is an essential precursor of the peptidoglycan cell wall (55, 56), and so is linked to both cell envelope stress and cellular nitrogen status. Deletion of *phoP* or *phoQ*, encoding a two-component regulator that responds to environmental stresses, including low pH and magnesium limitation, and is known to also positively regulate *glmY* (33), also led to a significant increase in polyP synthesis (Fig. 3B), consistent with this being the mechanism by which the Δ*glrR* mutation does so (Fig. 1B). PhoP is a global regulator that affects the expression and stability of many proteins (33), but is best known for its role in activating magnesium import (33, 57). High levels of polyP synthesis can cause magnesium starvation in *E. coli*, presumably by chelating the metal (23). However, deletion of the PhoP-regulated magnesium stress response genes *mgtA* and *mgtS* had no effect on polyP synthesis (Fig. 3B), and polyP-inducing nutrient shift did not activate the magnesium starvation response (Supplemental Fig. S1). PolyP-inducing nutrient limitation conditions did strongly downregulate *glmS* expression (Fig. 3C). Neither fructose-6-phosphate nor GlcN6P affected the activity of PPK *in vitro* (Fig. 3D).

**FIG 3.**
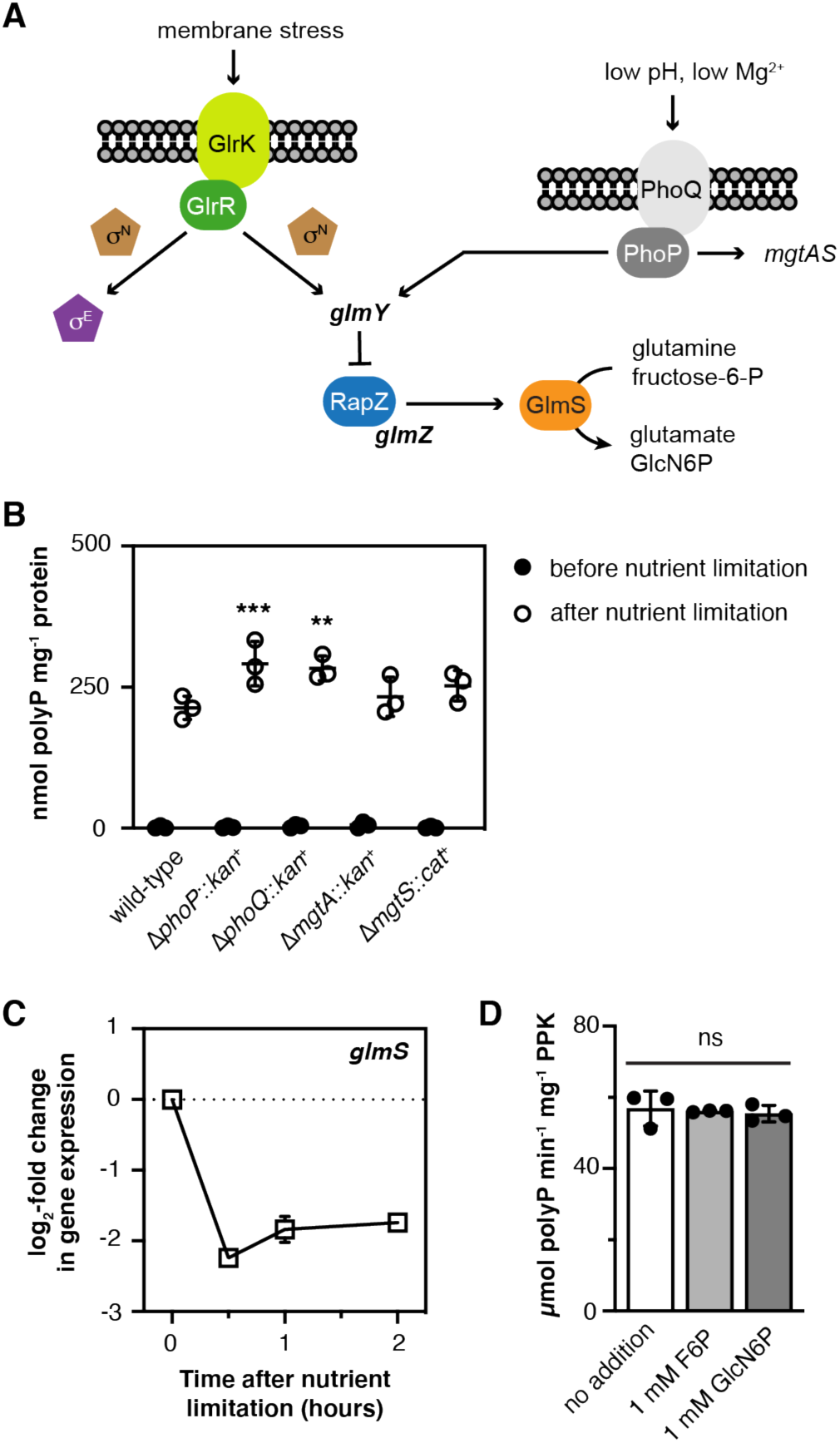
Effect of *glmY* and GlmS (glutamine--fructose-6-phosphate aminotransferase) regulation on polyP production. (*A*) Diagram of the GlrR- and PhoP-dependent pathways which regulate GlmS expression in *E. coli*. (*B*) MG1655 wild-type, Δ*phoP790*::*kan*^+^, Δ*phoQ789*::*kan*^+^, Δ*mgtA789*::*kan*^+^, or Δ*mgtS1000*::*cat^+^* were grown at 37°C to A_600_=0.2–0.4 in LB (black circles) and then shifted to minimal medium for 2 hours (white circles)(n=3, ± SD). (*C*) *E. coli* MG1655 wild-type was grown at 37°C to A_600_=0.2–0.4 in rich medium (LB) and then shifted to minimal medium (MOPS with no amino acids, 4 g l^-1^ glucose, 0.1 mM K_2_HPO_4_, 0.1 mM uracil) for 2 hours. qRT-PCR was used to measure fold changes in *glmS* transcript abundance at the indicated timepoints (n=3, ± SD), normalized to expression before stress treatment (t = 0 h). (*D*) *S*pecific activity of purified PPK in the presence of the indicated compounds (n=3, ±SD; one-way ANOVA, ns = not significant).

### PolyP regulation by RapZ and GlmS depends on growth conditions

Given the phenotypes of the Δ*glrR* and Δ*phoP* mutants (Figs. 1B, 3B), we hypothesized that deletion of *glmY* would increase the amount of polyP produced by *E. coli* and that either deletion of *rapZ* or *glmZ* or overexpression of *glmS* would decrease polyP production (Fig. 3A). However, under our normal nutrient limitation conditions, none of these had any effect on polyP synthesis (Figs. 4A,B). We wondered whether this was due to inadequate expression of *glmS* from an arabinose-inducible promoter in minimal medium containing glucose (58), so performed a nutrient shift into minimal medium containing arabinose as a sole carbon source (Fig. 4C). Under these conditions, polyP accumulation in the vector-only control was approximately half of that seen after nutrient shift into glucose, but *glmS* expression did significantly reduce polyP production. Nutrient shift from glutamine-supplemented rich medium into minimal glucose medium also resulted in modest, but statistically significant, defects in polyP synthesis in a Δ*rapZ* mutant and upon *glmS* overexpression (Figs. 4D,E). Nutrient shift from LB into minimal medium with glucosamine as a sole carbon source, expected to result in very high levels of intracellular glucosamine and low GlmS activity (59–61), resulted in significantly less polyP production than shift into glucose, and shift into glycerol, expected to result in low intracellular glucosamine and therefore high GlmS activity (51, 60) resulted in no detectable polyP production at all (Fig. 4F). The extent of polyP production was not affected by supplementation of the rich medium with glucosamine (Supplemental Fig. S2). These results reinforce the observation that growth conditions impact not only the extent of polyP synthesis, but the pathways involved in its regulation, and indicate that regulators of GlmS can, under certain circumstances, impact polyP production, with a general correspondence between higher polyP levels when GlmS levels are low and *vice versa*, but can not fully explain the mechanism by which this occurs.

**FIG 4.**
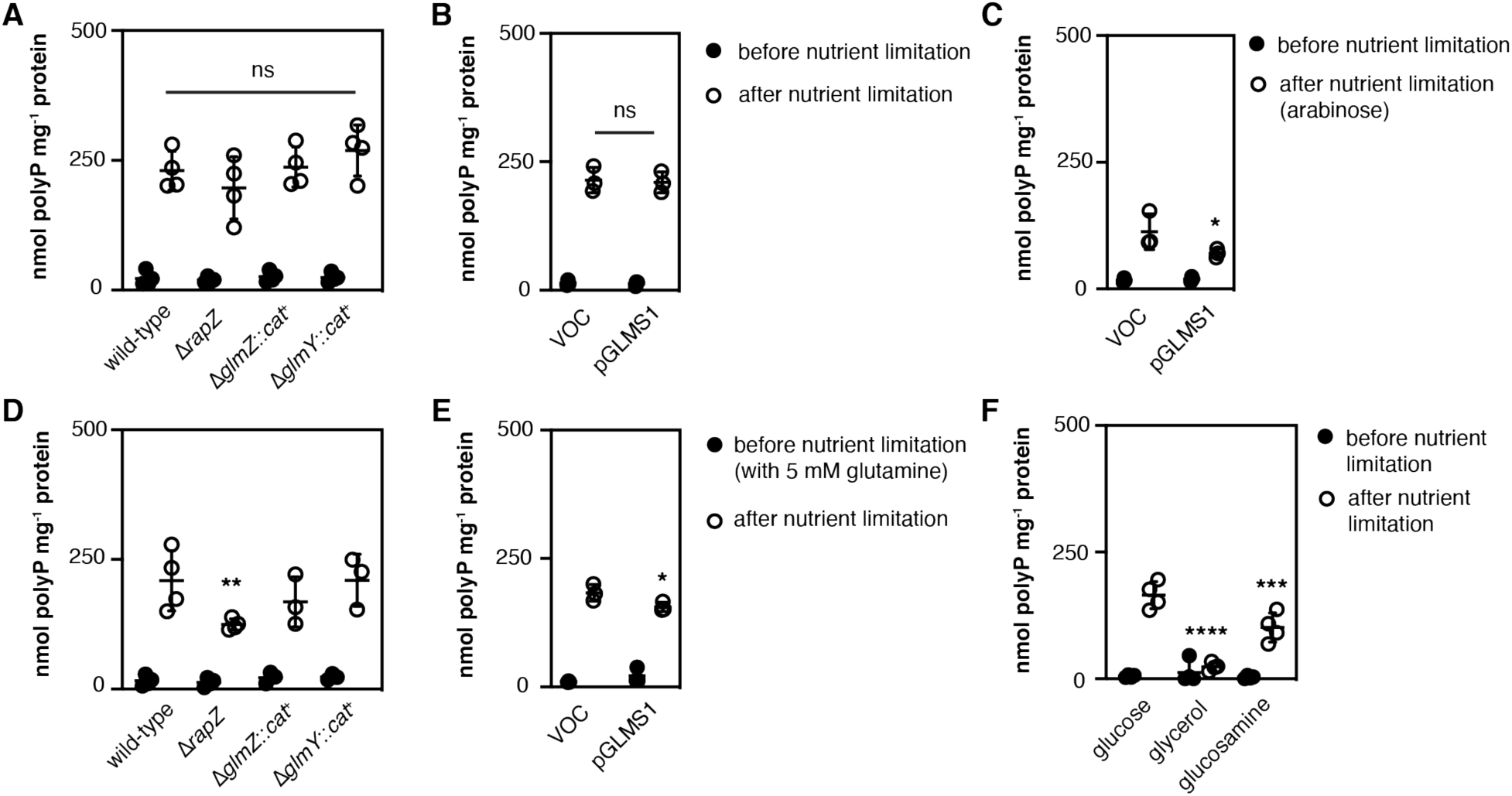
PolyP regulation by RapZ and GlmS depends on growth conditions. (*A*) *E. coli* MG1655 wild-type, Δ*rapZ733*, Δ*glmZ1000*::*cat^+^*, or Δ*glmY1000*::*cat^+^* were grown at 37°C to A_600_=0.2–0.4 in LB (black circles) and then shifted to minimal medium for 2 hours (white circles)(n=4, ± SD). (*B*) *E. coli* MG1655 wild-type containing plasmids pBAD18 (VOC) or pGLMS1 (*glmS*^+^) were grown at 37°C to A_600_=0.2–0.4 in LB containing 2 g l^-1^ arabinose (black circles) and then shifted to minimal medium supplemented with 2 g l^-1^ arabinose for 2 hours (white circles)(n=3, ± SD). (*C*) *E. coli* MG1655 wild-type containing plasmids pBAD18 or pGLMS1 were grown at 37°C to A_600_=0.2–0.4 in LB containing 2 g l^-1^ arabinose (black circles) and then shifted to minimal medium containing 4 g l^-1^ arabinose as a sole carbon source for 2 hours (white circles)(n=3, ± SD). (*D*) *E. coli* MG1655 wild-type, Δ*rapZ733*, Δ*glmZ1000*::*cat^+^*, or Δ*glmY1000*::*cat^+^* were grown at 37°C to A_600_=0.2–0.4 in LBQ (black circles) and then shifted to minimal medium containing for 2 hours (white circles)(n=3-4, ± SD). (*E*) *E. coli* MG1655 wild-type containing plasmids pBAD18 or pGLMS1 were grown at 37°C to A_600_=0.2–0.4 in LBQ containing 2 g l^-1^ arabinose (black circles) and then shifted to minimal medium supplemented with 2 g l^-1^ arabinose for 2 hours (white circles)(n=3, ± SD). (*F*) *E. coli* MG1655 wild-type was grown at 37°C to A_600_=0.2–0.4 in LB (black circles) and then shifted to minimal media containing either 4 g l^-1^ glucose, 8 g l^-1^ glycerol, or 4 g l^-1^ glucosamine as sole carbon sources for 2 hours (white circles)(n=3-4, ± SD). PolyP concentrations are in terms of individual phosphate monomers. Asterisks indicate polyP levels significantly different from those of the wild-type control for a given experiment unless otherwise indicated (two-way repeated measures ANOVA with Holm-Sidak’s multiple comparisons test, ns = not significant * = P<0.05, ** = P<0.01, *** = P<0.001, **** = P<0.0001).

### PtsN positively regulates polyP synthesis regardless of its phosphorylation state or the presence of glutamine, but does not do so by interacting directly with PPK

The nitrogen phosphotransferase system (PTS^Ntr^) is a regulatory cascade that responds to nitrogen limitation and regulates the activity of multiple proteins in *Enterobacteriacea*, including both GlmS and PhoP (62–67)(Fig. 5A). The genes encoding PtsN (also known as EIIA^Ntr^) and NPr, homologs of the EIIA and HPr proteins of the well-characterized carbon PTS (68), are encoded in the *rpoN* operon (34). PtsP (also known as EI^Ntr^), which is homologous to EI of the carbon PTS (34, 68), responds to both glutamine and *α*-ketoglutarate as signals of cellular nitrogen limitation by autophosphorylation (37, 69), ultimately resulting in phosphorylation of NPr and PtsN (70). Both NPr and PtsN interact with and regulate the activity of a variety of proteins, typically depending on their phosphorylation states (62–67). Based on these facts and our results showing roles for PhoP and GlmS in polyP regulation we hypothesized that the PTS^NTr^ might form part of the link between nitrogen limitation and polyP synthesis.

**FIG 5.**
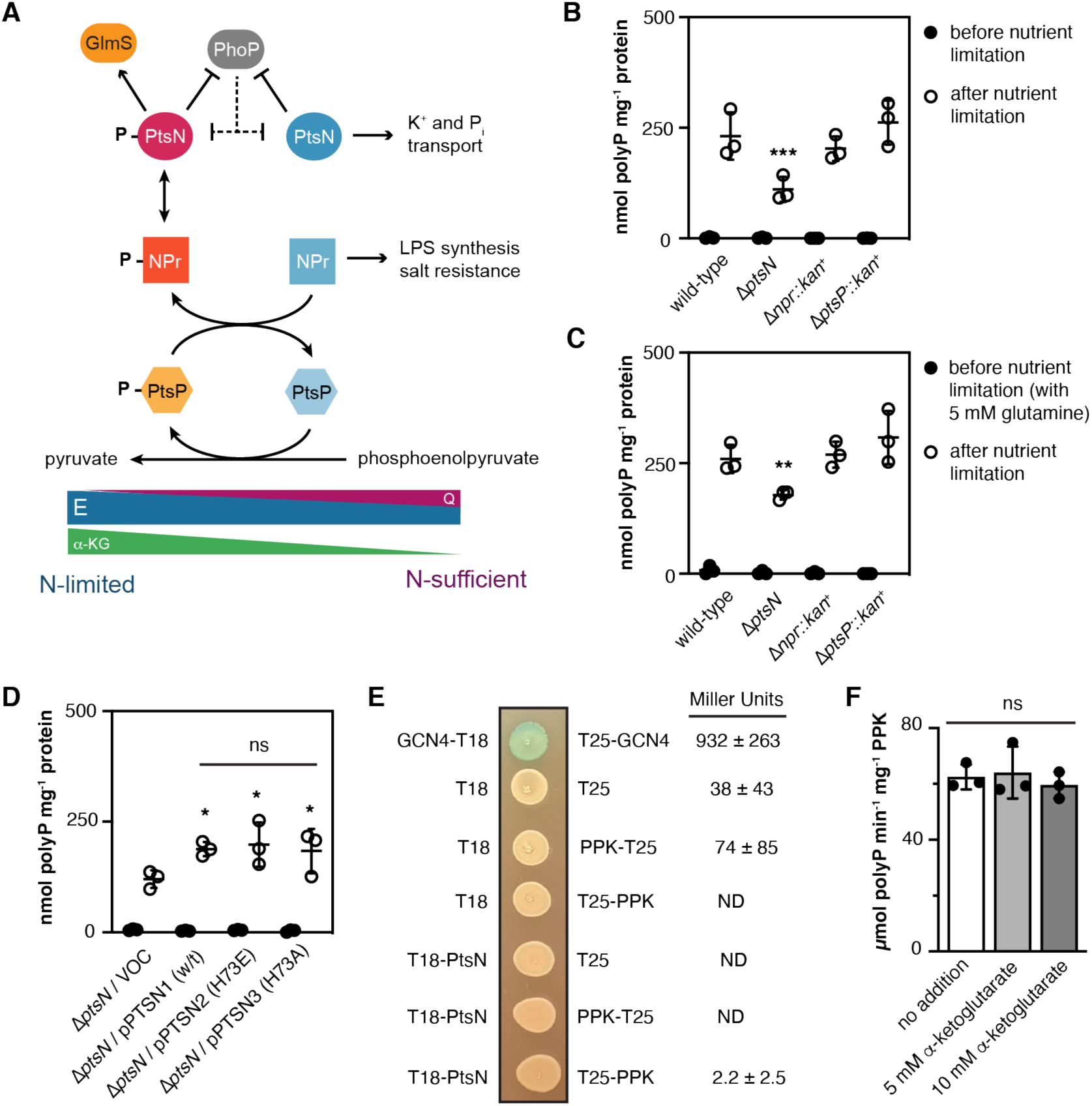
PtsN positively regulates polyP synthesis regardless of its phosphorylation state or the presence of glutamine, but does not do so by interacting directly with PPK. (*A*) Diagram of the nitrogen phosphotransferase system of *E. coli*. (*B,C*) MG1655 wild-type, Δ*ptsN732*, Δ*npr-734*::*kan*^+^, or Δ*ptsP753*::*kan*^+^ were grown at 37°C to A_600_=0.2–0.4 in LB (panel A) or LBQ (panel B) (black circles) and then shifted to minimal medium for 2 hours (white circles)(n=3, ± SD). (*D*) MG1655 Δ*ptsN732* containing the indicated plasmids was grown at 37°C to A_600_=0.2–0.4 in LB containing 2 g l^-1^ arabinose (black circles) and then shifted to minimal medium supplemented with 2 g l^-1^ arabinose for 2 hours (white circles)(n=3, ± SD). VOC is pBAD18. PolyP concentrations are in terms of individual phosphate monomers. Asterisks indicate polyP levels significantly different from those of the wild-type or Δ*ptsN* / VOC control for a given experiment unless otherwise indicated (two-way repeated measures ANOVA with Holm-Sidak’s multiple comparisons test, ns = not significant * = P<0.05, ** = P<0.01, *** = P<0.001, **** = P<0.0001). (*E*) *E. coli* BTH101 (*cya*^-^) containing plasmids expressing the indicated protein fusions were grown overnight in LB and either spotted on LB medium containing 0.5 mM IPTG and 40 µg ml^-1^ X-Gal or lysed for quantitative assay of β-galactosidase activity (n=3, ± SD; ND = not detectable). (*F*) *S*pecific activity of purified PPK in the presence of the indicated compounds (n=3, ±SD; one-way ANOVA, ns = not significant).

Mutants lacking *ptsN* were defective in polyP synthesis (Fig. 5B). However, this defect was not affected by glutamine supplementation and was not seen in Δ*npr* or Δ*ptsP* mutants (Figs. 5B,C), suggesting that phosphorylation was not important for this phenotype. Indeed, both the non-phosphorylatable PtsN^H73A^ variant and the phosphorylated form-mimicking PtsN^H73E^ variant (71, 72) complemented the polyP defect of a Δ*ptsN* mutant as well as did wild-type PtsN (Fig. 5D). PtsN regulates other proteins by direct physical interaction (62–67), so we used a bacterial two-hybrid assay (73) to test whether PtsN interacts with PPK *in vivo*, and found that it does not (Fig. 5E). PPK activity is also not affected by *α*-ketoglutarate *in vitro* (Fig. 5F), eliminating this metabolite as a possible allosteric regulator of polyP production.

### Effect of polyP-inducing nutrient limitation on PtsN levels *in vivo*

The only reported example of a protein in *Enterobacteria* that interacts with both the phosphorylated and dephosphorylated forms of PtsN is PhoP (33, 62). In *Salmonella*, PtsN inhibits PhoP binding to DNA, and in turn, PhoP regulates the proteolytic degradation of PtsN, leading to a decrease in PtsN protein concentration under PhoP-activating conditions (Fig. 5A). Both abundance and phosphorylation of PtsN are therefore potential variables in any PtsN-dependent regulatory system. PtsN abundance in *Salmonella* is also regulated in response to carbon source availability (63). We constructed strains encoding chromosomal fusions of the 3xFLAG epitope tag to the C-terminus of PtsN (62) to allow us to determine whether our polyP-inducing stress conditions led to changes in PtsN abundance in *E. coli*. In wild-type and Δ*phoP* strains (Figs. 6A,B), there was no significant change in PtsN abundance after nutrient limitation, consistent with the apparent lack of PhoP induction under these conditions (Supplemental Fig. S1). There was a significant increase in PtsN abundance two hours after nutrient limitation in a Δ*rpoN* mutant (Fig. 6C), but this increase did not correlate with polyP accumulation, which is lower in this strain (Fig. 1A). Based on these results, the regulation of polyP by PtsN does not appear to depend on PtsN abundance.

**FIG 6.**
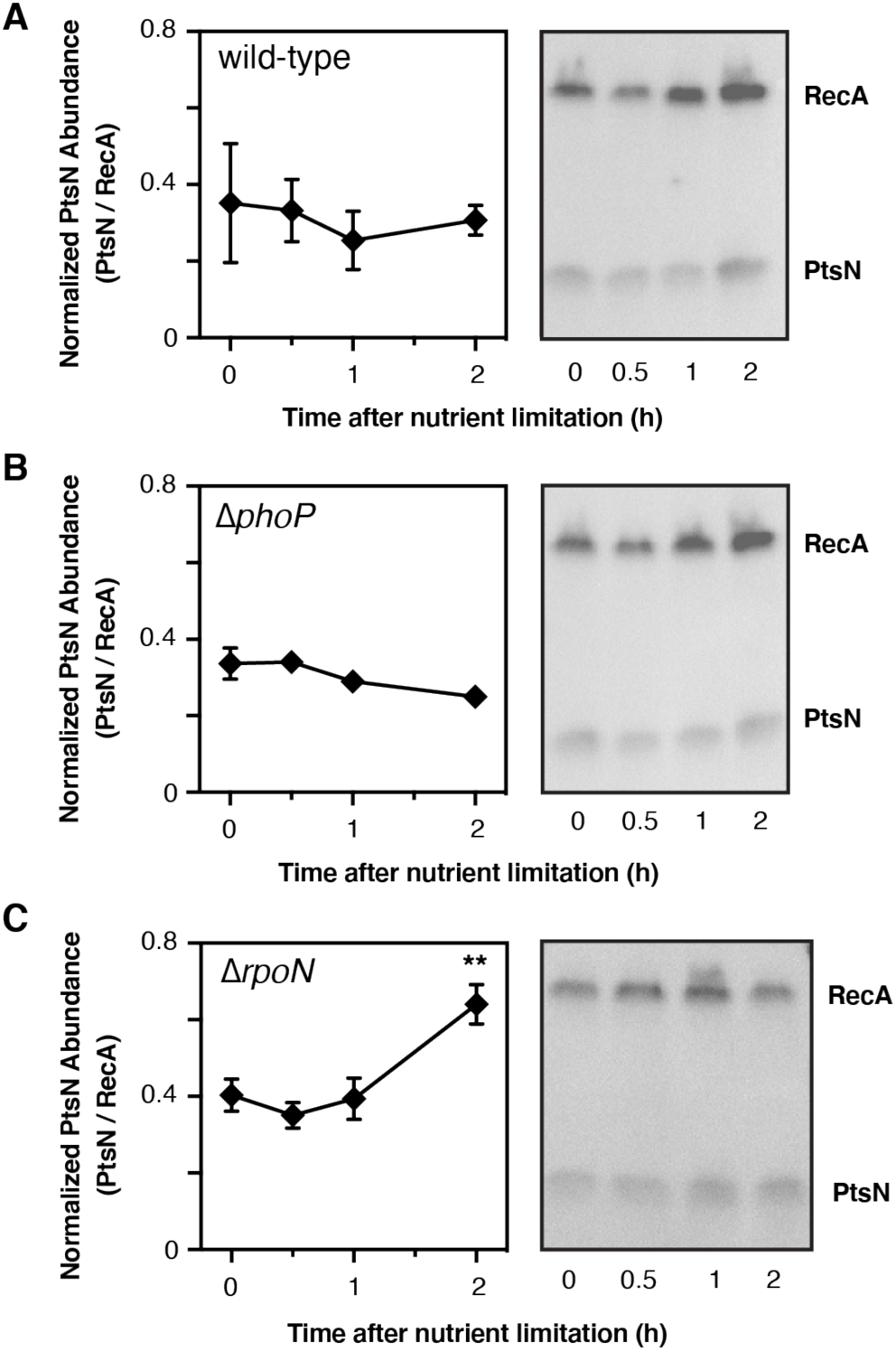
Effect of polyP-inducing nutrient limitation on PtsN levels *in vivo*. *E. coli* MG1655 (*A*) *ptsN*-3xFLAG, (*B*) Δ*phoP790*::*kan*^+^ *ptsN*-3xFLAG, or (*C*) Δ*rpoN730 ptsN*-3xFLAG strains were grown at 37°C to A_600_=0.2–0.4 in LB and then shifted to minimal medium for 2 hours. At the indicated timepoints, protein samples (n=3, ±SD) were collected and immunoblotted to quantify PtsN-3xFLAG and RecA. Representative blots are shown. Full gels are shown in Supplemental Fig. S3. Asterisks indicate normalized PtsN levels significantly different from those of the wild-type at the indicated timepoint (two-way repeated measures ANOVA with Holm-Sidak’s multiple comparisons test, ** = P<0.01).

### PtsN acts downstream of RpoN, upstream of DksA, and independently of RpoE in polyP regulation

Finally, we used complementation analysis to examine the relationships between the different regulators of polyP production we have identified. PtsN, RpoN, DksA, and RpoE are all positive regulators of polyP production (Figs. 1, 5)(22, 24). Expressing RpoN from a plasmid does not increase polyP synthesis in a Δ*ptsN* mutant (Fig. 7A), but expressing either DksA or RpoE does (Fig. 7B). In contrast, expressing PtsN rescues polyP production in Δ*rpoN* and Δ*rpoE* mutants, but not in a Δ*dksA* mutant (Figs. 7C,D,E). This suggests a pathway in which, in response to nutrient limitation stress, RpoN positively regulates PtsN, which in turn positively regulates DksA, increasing polyP production. We do not, at this time, know which of these activation steps are direct and which are indirect or the mechanism by which DksA activates polyP production (22, 24). The case of RpoE is more complicated. Since the defects of Δ*rpoE* and Δ*ptsN* mutants can each be rescued by expression of the other gene (Figs. 7B,E), the simplest interpretation is that they regulate polyP production by independent mechanisms. However, *ptsN* is both a member of the RpoE regulon (74) and a multicopy suppressor of the conditionally lethal phenotype of a Δ*rpoE* mutant (75), so these results must be interpreted cautiously. Expression of RpoN rescues the polyP defect of a Δ*rpoE* mutant (22), DksA is a positive regulator of RpoE-dependent transcription for some genes (76), and there are known interactions between RpoN and RpoE in response to combined nitrogen and carbon starvation stress conditions (77). We think it is unlikely, therefore, that these regulators are operating completely independently in control of polyP production, but do not yet have enough data to speculate on their exact relationship.

**FIG 7.**
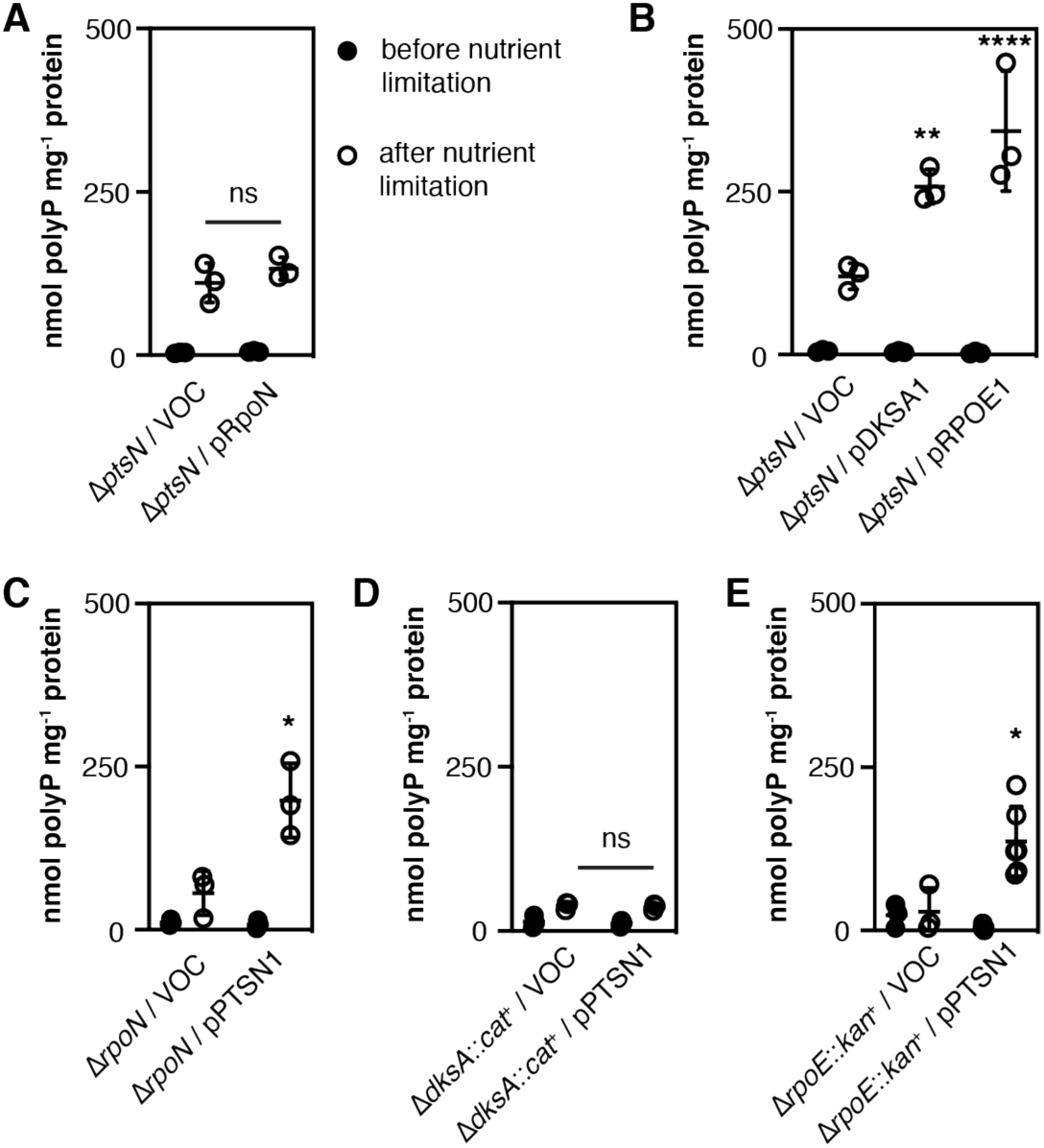
PtsN acts downstream of RpoN, upstream of DksA, and independently of RpoE in polyP regulation. MG1655 Δ*ptsN732* (panels *A, B*) Δ*rpoN730* (panel *C*), Δ*dksA1000*::*cat*^+^ (panel *D*), or Δ*rpoE1000*::*kan*^+^ (panel *E*) strains containing the indicated plasmids were grown at 37°C to A_600_=0.2–0.4 in LB containing 2 g l^-1^ arabinose (black circles) and then shifted to minimal medium supplemented with 2 g l^-1^ arabinose for 2 hours (white circles)(n=3, ± SD). VOC for panel A is pBAD24 and for panels B-E is pBAD18, and both the LB and minimal medium contained 10 µg ml^-1^ erythromycin for the experiment shown in panel E. PolyP concentrations are in terms of individual phosphate monomers. Asterisks indicate polyP levels significantly different from those of the VOC control for a given experiment (two-way repeated measures ANOVA with Holm-Sidak’s multiple comparisons test, ns = not significant * = P<0.05, ** = P<0.01, *** = P<0.001, **** = P<0.0001).

## DISCUSSION

It is perhaps unsurprising that the regulation of polyP synthesis is complex, given the ancient evolutionary roots of polyP (3, 6, 78, 79) and the intricacy of the regulatory networks for other general stress response pathways in bacteria. In *E. coli*, for example, there are at least 20 regulators of the stress-responsive sigma factor RpoS known at time of writing, acting at the transcriptional, post-transcriptional, and post-translational levels (27, 80, 81). Envelope stress responses are equally complex and involve a variety of interacting and overlapping pathways and regulons, the details of which are still not fully understood (53, 82).

The experiments presented in this paper were intended to identify the gene or genes regulated by RpoN that contribute to polyP production (16, 22), and while we did successfully identify roles for the RpoN-related proteins GlnG, GlrR, and PtsN in modulating polyP production (Figs. 1, 5), we did not find a simple RpoN-dependent activator of polyP production and uncovered unexpected impacts of PhoPQ, RapZ, and GlmS (Figs. 3, 4). Some of these regulators only impacted polyP production under specific growth conditions. Adding glutamine to the rich media before nutrient limitation makes RpoN and GlnG unneccessary for polyP production (Fig. 2F), but makes RapZ and GlmS expression significant regulators (Fig. 4D,E). The carbon source in the minimal medium also affects both polyP production and the impact of GlmS expression on that production (Fig. 4C,F). Based on this, one important conclusion we can draw from our results is that, while shift from rich medium to minimal medium is in general a strong signal inducing polyP production (16, 24), the composition of both the rich and minimal media impacts not only the extent of polyP production, but also the pathways involved in regulating that production. This is not, in hindsight, completely surprising, but does mean that growth in LB medium (50) can not simply be considered a “non-stress” condition that contrasts with “stressful” nutrient limitation, and that in order to fully understand polyP regulation we will also need to consider the cell’s physiological state under conditions when it is not producing detectable amounts of polyP.

The exact function of the PTS^Ntr^ has been debated for some time (28, 34), but it is clear from our results that PtsN has a phosphorylation-independent positive effect on polyP synthesis (Figs. 5, 7). What is less clear is how this occurs. PtsN does not appear to interact directly with PPK *in vivo* (Fig. 5E). Overexpression of PtsN is known to generally reduce cell envelope stress in *E. coli*, although the mechanism by which it does so is not well understood (75). Known targets of PtsN regulation include proteins involved in phosphate transport (83), potassium transport (84–86), GlcN6P synthesis (64), and environmental sensing (62). While phosphate transport is certainly important for polyP synthesis (16, 19, 87, 88) and PtsN-dependent changes in potassium levels are known to impact sigma factor specificity (85), which may also play a role in polyP regulation (22), both of these phenotypes are dependent on the phosphorylation state of PtsN, as is the interaction between PtsN and GlmS (64). While this manuscript was in preparation, Gravina *et al*. (89) reported the identification of multiple new PtsN interaction candidates in *E. coli*, most of which also appear to be phosphorylation-state specific. Fortuitiously, we have already tested many of these for potential roles in polyP accumulation (22), including proteins involved in flagellar motility and glycerol metabolism, and found that those pathways have minimal effects on polyP accumulation. However, there are additional candidates, including proteins of unknown function (*e.g.* YeaG, YcgR, and YcjN) and enzymes of central metabolism (*e.g.* AceAB, PpsA, SucC), that remain to be tested.

Nevertheless, with the information in hand, we identified the interaction between PtsN and the transcription factor PhoP (62) as the most likely candidate for a phosphorylation-independent mechanism for PtsN’s effect on polyP accumulation. PhoP is a global regulator that responds to magnesium limitation, low pH, antimicrobial peptides, hyperosmotic stress, periplasmic redox state, and other environmental conditions (33). PtsN inhibits PhoP’s DNA binding affinity, thereby potentially impacting transcription of the PhoP regulon and of genes controlled by regulators that are part of that regulon (*e.g.* RstA, MgrR,or IraM)(33, 62, 90). However, PhoP also has large-scale post-translational effects on the proteome by regulating the activity of proteases (33, 91, 92), and PtsN is degraded by the Lon protease under PhoP-activating conditions in a PhoP-dependent feedback loop (62). While PtsN abundance was not regulated in response to nutrient limitation in wild-type or Δ*phoP* strains (Figs. 6A,B), PtsN abundance did increase at late time points in a Δ*rpoN* mutant (Fig. 6C). It is tempting to speculate that this mutant was upregulating PtsN in an attempt to compensate for the loss of RpoN (Fig. 7C), but additional experiments will be needed to test this hypothesis. The increase in polyP production in *phoPQ* mutants (Fig. 3B) is consistent with the hypothesis that PtsN acts in concert with PhoP to regulate polyP production, but it is also possible that there is another PtsN interaction partner involved. Which member(s) of the PhoP regulon besides PtsN might affect polyP production is also unknown.

It is important to note that none of the regulators we identified in this paper are absolutely required for induction of polyP synthesis after stress. Some impact polyP production positively (PtsN, GlnG) and some negatively (GlrR, PhoPQ), but every mutant we tested was still able to respond to nutrient limitation stress by increasing polyP production to some extent. This probably indicates that there are redundant mechanisms by which stress signals impact PPK and PPX activity. We know this to be the case for PPX, which is inhibited by both (p)ppGpp and hypochlorous acid-driven oxidation (17, 20). The mechanism(s) by which PPK activity is controlled remain unknown, although the stress-responsive accumulation of polyP in Δ*ppx* mutant strains indicates that such a mechanism must exist (17, 24). Our *in vitro* results (Figs. 2G, 3D, 5F) do show that PPK itself is not allosterically regulated by a set of common metabolites whose levels change under nitrogen limitation conditions (28, 37).

What is clear from our results (Figs. 1, 3, 4, 5) is that multiple pathways that influence cell envelope stress or synthesis impact polyP synthesis in *E. coli*. Without knowing the mechanism by which PPK activity itself is modulated, it is difficult to determine whether these pathways directly regulate polyP synthesis or, alternatively, whether polyP synthesis responds to the changes in cell envelope homeostasis that result from disruption of those regulatory networks (28-30, 32, 33). Based on the data in this and our previous papers (17, 22–24), on balance, we favor the second hypothesis, but are working to clarify this question. PPK is a peripheral membrane protein (93), so it is not impossible that PPK itself is sensitive to changes in the cytoplasmic membrane *in vivo*. If this is the case, it will be difficult to recapitulate PPK regulation *in vitro*. Regardless, our results illustrate previously unknown connections among a variety of well-conserved environmental stress response pathways and show that even as well-studied an organism as *E. coli* still has plenty of capacity to surprise us and confound our expectations.

## MATERIALS AND METHODS

### Bacterial strains, growth conditions, and molecular methods

All strains and plasmids used in this study are listed in Table 1. We carried out DNA manipulations by standard methods (94, 95) in the *E. coli* cloning strain DH5*α* (Invitrogen) and grew *E. coli* at 37°C in Lysogeny Broth (LB)(96) containing 5 g l^-1^ NaCl and, where indicated, 5 mM *L*-glutamine (LBQ) or 4 g l^-1^ glucosamine (LBGlcN). We prepared fresh glutamine and glucosamine stock solutions each day. We added antibiotics when appropriate: ampicillin (100 µg ml^-1^), chloramphenicol (17 or 35 µg ml^- 1^), gentamycin (30 µg ml^-1^), kanamycin (25 or 50 µg ml^-1^), or spectinomycin (50 µg ml^-1^). We constructed, maintained, and tested all *rpoE* mutant strains in media containing 10 µg ml^-1^ erythromycin (97).

**TABLE 1.**
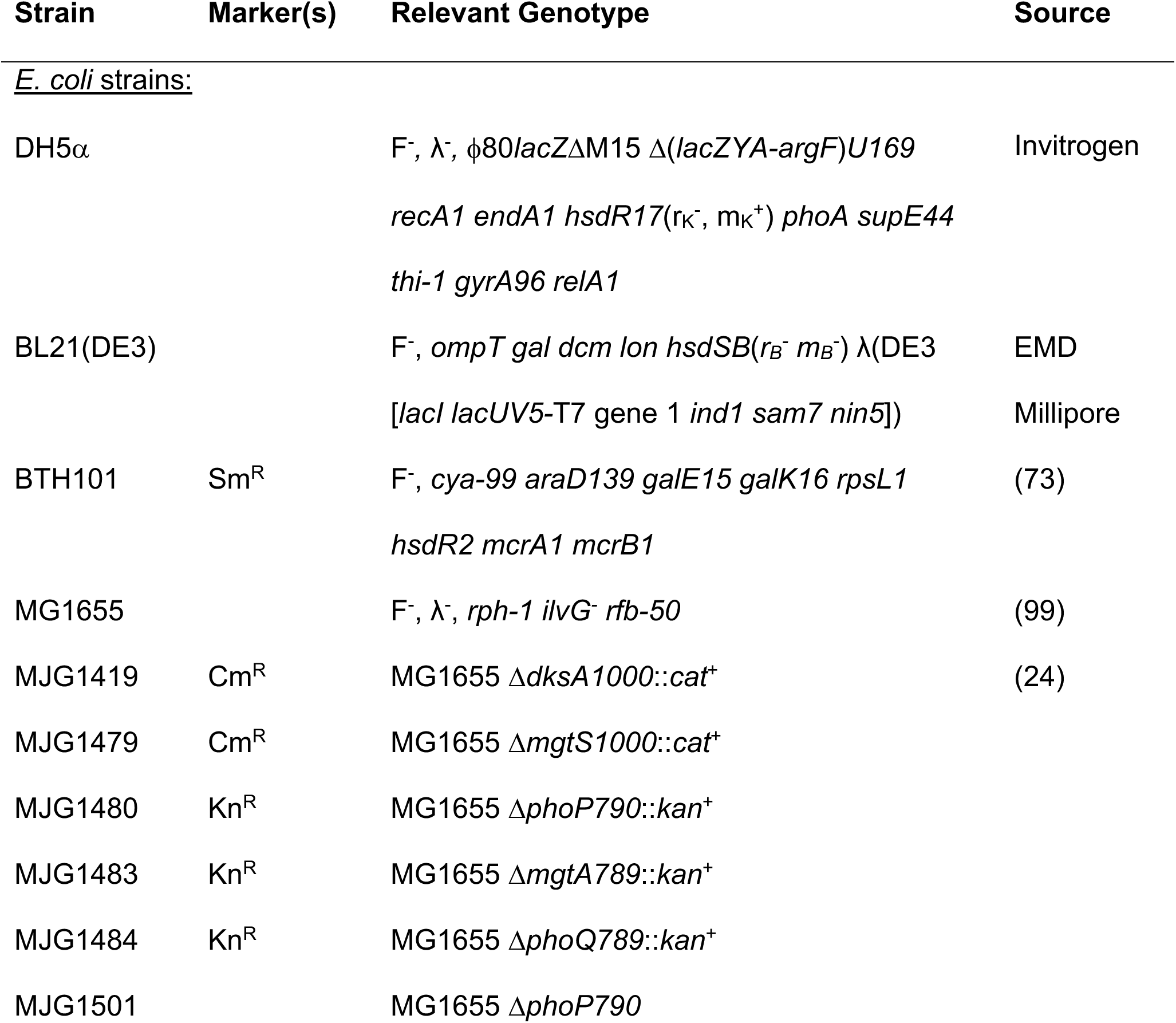
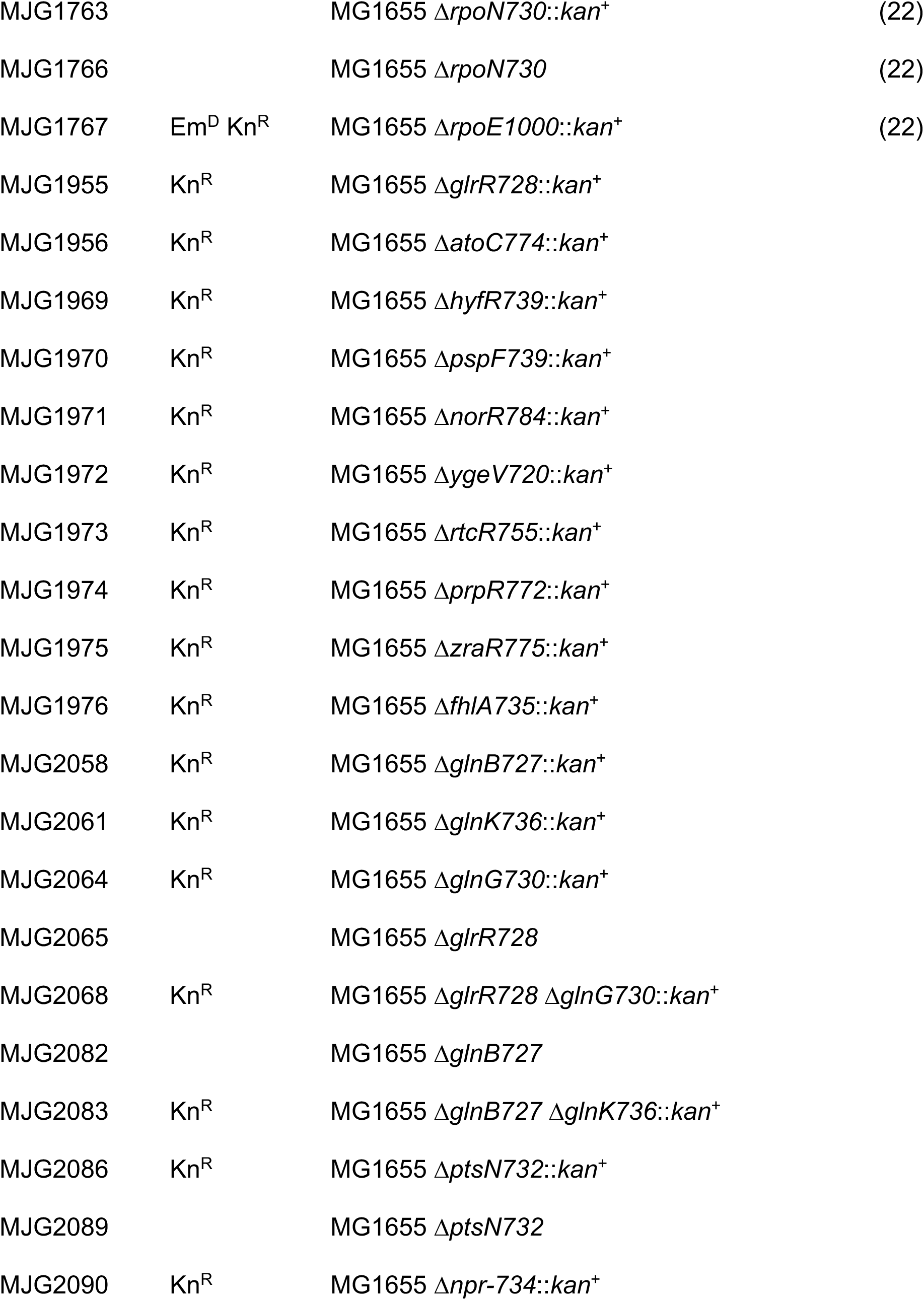
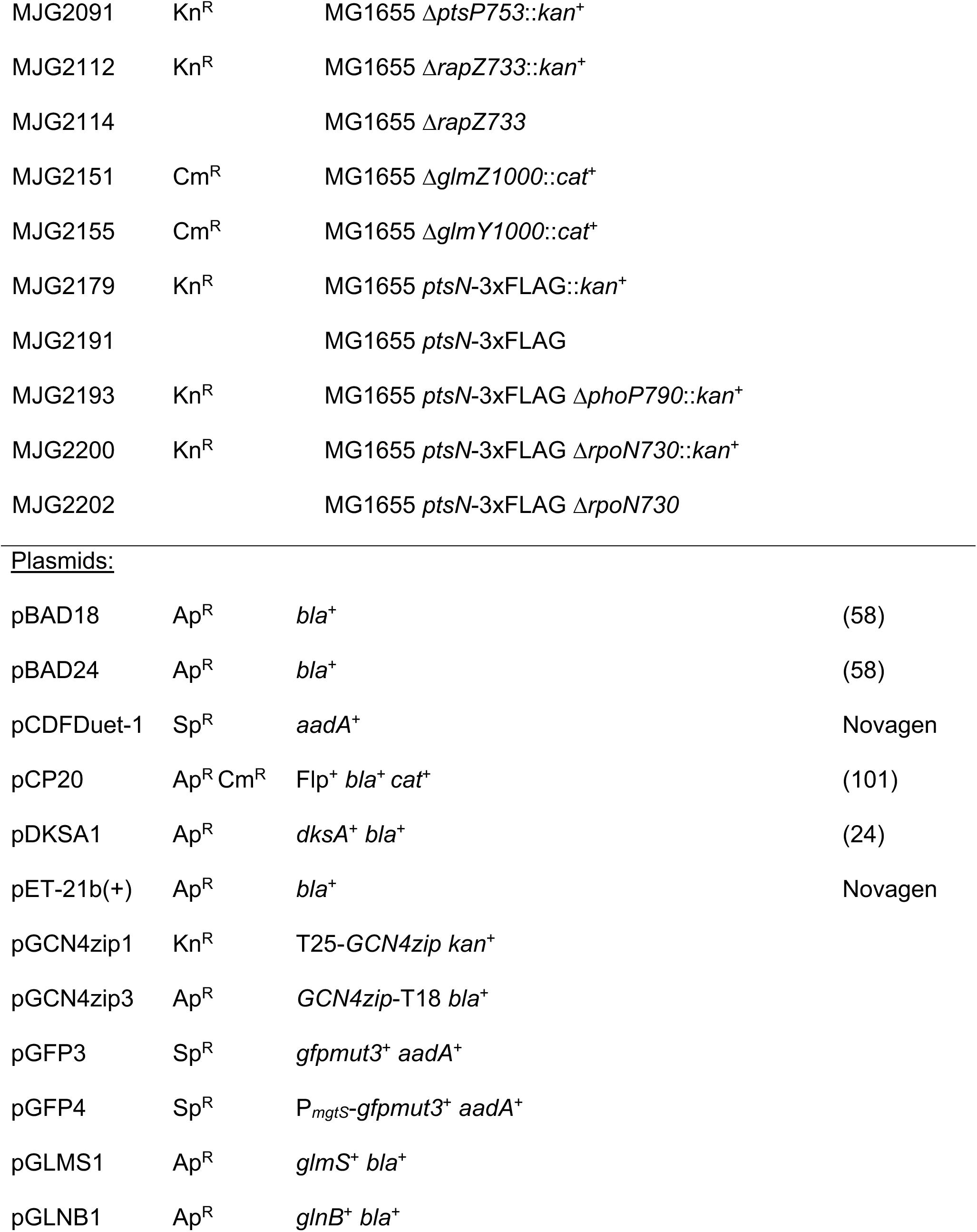
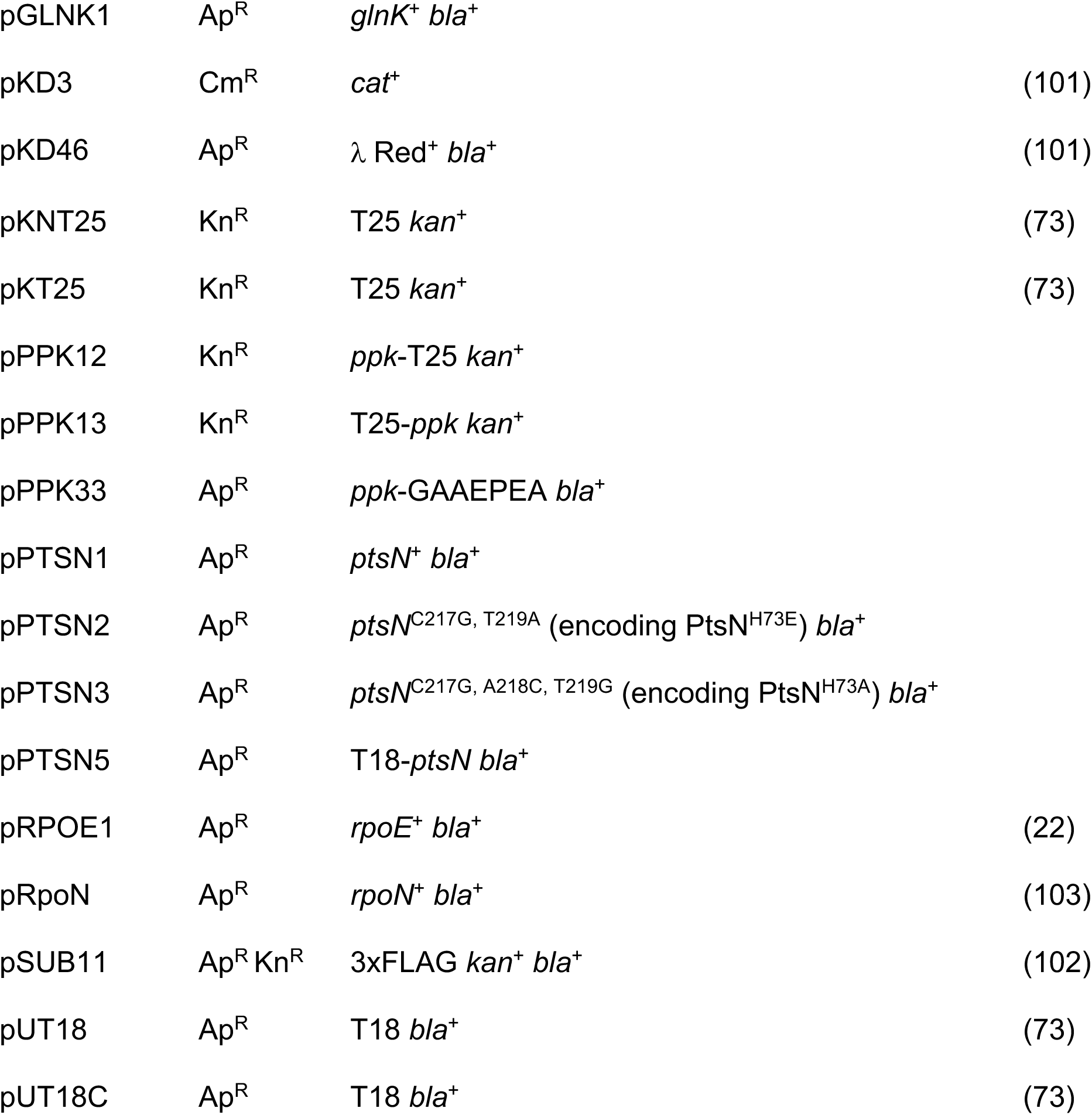
Strains and plasmids used in this study. Unless otherwise indicated, all strains and plasmids were generated in the course of this work. Abbreviations: Ap^R^, ampicillin resistance; Cm^R^, chloramphenicol resistance; Em^D^, erythromycin dependance; Gm^R^, gentamycin resistance; Kn^R^, kanamycin resistance; Sp^R^, spectinomycin resistance; Sm^R^, streptomycin resistance.

### Databases and primer design

We obtained information about *E. coli* genes, proteins, and regulatory networks from the Integrated Microbial Genomes database (98), EcoCyc (27), and RegulonDB (45). We designed PCR and sequencing primers with Web Primer (www.candidagenome.org/cgi-bin/compute/web-primer) or SnapGene version 5.3.2 (Insightful Science), and mutagenic primers with PrimerX (www.bioinformatics.org/primerx/index.htm). We designed all primers used for qPCR with Primer Quest (www.idtdna.com; parameter set “qPCR 2 primers intercalating dyes” for qRT-PCR primer design) and confirmed specificity and amplification efficiencies for each primer pair of close to 1. These primers are listed in Table 2.

**TABLE 2.**
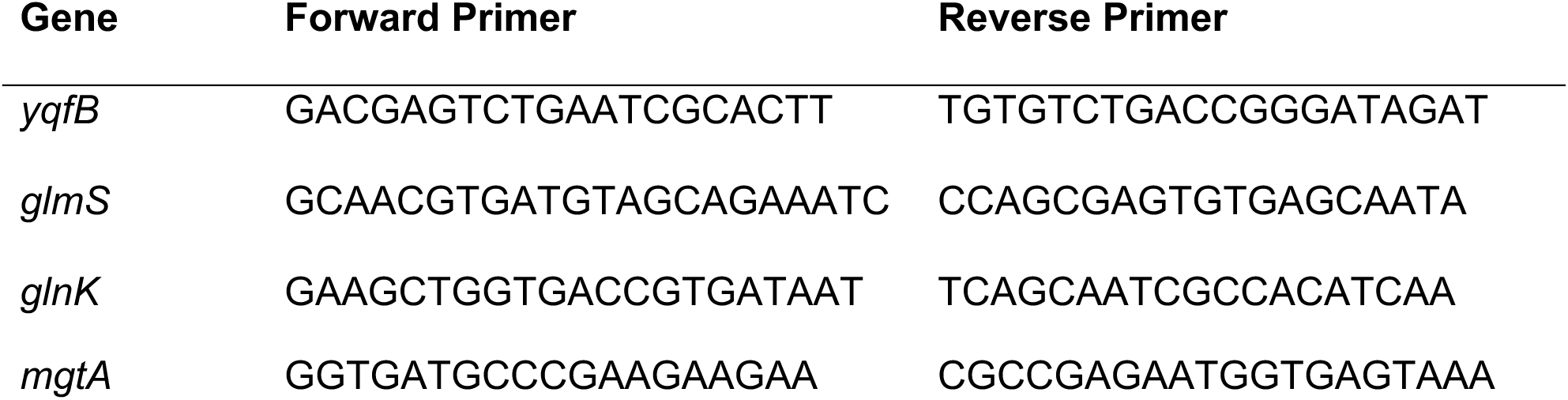
Primers used for quantitative RT-PCR

### Strain construction

Unless otherwise indicated, all *E. coli* strains were derivatives of wild-type strain MG1655 (F^-^, *rph-1 ilvG^-^ rfb-50*)(99), and we confirmed all chromosomal mutations by PCR.

We used P1*vir* transduction (24, 100) to move gene knockout alleles from the Keio collection (43) into MG1655, generating strains MJG1480 (Δ*phoP790*::*kan*^+^), MJG1483 (Δ*mgtA789*::*kan*^+^), MJG1484 (Δ*phoQ789*::*kan*^+^), MJG1955 (Δ*glrR728*::*kan*^+^), MJG1956 (Δ*atoC774*::*kan*^+^), MJG1969 (Δ*hyfR739*::*kan*^+^), MJG1970 (Δ*pspF739*::*kan*^+^), MJG1971 (Δ*norR784*::*kan*^+^), MJG1972 (Δ*ygeV720*::*kan*^+^), MJG1973 (Δ*rtcR755*::*kan*^+^), MJG1974 (Δ*prpR772*::*kan*^+^), MJG1975 (Δ*zraR775*::*kan*^+^), MJG1976 (Δ*fhlA735*::*kan*^+^), MJG2058 (Δ*glnB727*::*kan*^+^), MJG2061 (Δ*glnK736*::*kan*^+^), MJG2064 (Δ*glnG730*::*kan*^+^), MJG2086 (Δ*ptsN732*::*kan*^+^), MJG2090 (Δ*npr-734*::*kan*^+^), MJG2091 (Δ*ptsP753*::*kan*^+^), and MJG2112 (Δ*rapZ733*::*kan*^+^). We resolved the insertions (101) in MJG1480, MJG1955, MJG2058, MJG2086, and MJG2112 to give strains MJG1501 (Δ*phoP790*), MJG2065 (Δ*glrR728*), MJG2082 (Δ*glnB727*), MJG2089 (Δ*ptsN732*), and MJG2114 (Δ*rapZ733*), then transduced MJG2065 and MJG2082 with Δ*glnG730*::*kan*^+^ and Δ*glnK736*::*kan*^+^, respectively, to yield strains MJG2068 (Δ*glrR728* Δ*glnG730*::*kan*^+^) and MJG2083 (Δ*glnB727* Δ*glnK736*::*kan*^+^). Strains lacking both *glnB* and *glnK* are glutamine auxotrophs (39) and were constructed and maintained on LBQ.

We replaced the *mgtS*, *glmZ,* and *glmY* genes of strain MG1655 with pKD3-derived chloramphenicol resistance cassettes by recombineering (101), using primers 5’ AAT TAA GGT AAG CGA GGA AAC ACA CCA CAC CAT AAA CGG AGG CAA ATA ATG GTG TAG GCT GGA GCT GCT TC 3’ and 5’ ACA CAA CTG TAA CAA GGG GCC GGT TAG GTG AGG GAT TAT CTC CGT TCA TTA CAT ATG AAT ATC CTC CTT AG 3’, 5’ AAG TGT TAA GGG ATG TTA TTT CCC GAT TCT CTG TGG CAT AAT AAA CGA GTA GTG TAG GCT GGA GCT GCT TC 3’ and 5’ CTT CCT GAT ACA TAA AAA AAC GCC TGC TCT TAT TAC GGA GCA GGC GTT AAA CAT ATG AAT ATC CTC CTT AG 3’, or 5’ TTA CCA AAC TAT TTT CTT TAT TGG CAC AGT TAC TGC ATA ATA GTA ACC AGT GTG TAG GCT GGA GCT GCT TC 3’ and 5’ TCG TCA GAC GCG AAT AGC CTG ATG CTA ACC GAG GGG AAG TTC AGA TAC AAC CAT ATG AAT ATC CTC CTT AG 3’, to yield strains MJG1479 (Δ*mgtS1000*::*cat*^+^), MJG2151 (Δ*glmZ1000*::*cat*^+^), and MJG2155 (Δ*glmY1000*::*cat*^+^).

We fused a 3xFLAG tag to the C-terminus of the chromosomal *ptsN* gene by recombineering (102). We amplified the 3xFLAG sequence and kanamycin resistance cassette from plasmid pSUB11 (102) with primers 5’ GAA GAG CTG TAT CAA ATC ATT ACG GAT ACC GAA GGT ACT CCG GAT GAA GCG GAC TAC AAA GAC CAT GAC GG 3’ and 5’ TAC CAT GTA CTG TTT CTC CTC ACA ACG TCT AAA AGA GAC ATT ACC GAA TAA CAT ATG AAT ATC CTC CTT AG 3’ and electroporated the resulting PCR product into MG1655 expressing *λ* Red recombinase from plasmid pKD46 (101), generating strain MJG2179 (*ptsN*-3xFLAG *kan*^+^). We then resolved the kanamycin resistance cassette in MJG2179 with plasmid pCP20 (101) to yield strain MJG2191 (*ptsN*-3xFLAG). We used P1*vir* transduction (24, 100) to move the Δ*phoP790*::*kan*^+^ allele from the Keio collection (43) into MJG2191, generating strain MJG2193 (*ptsN*-3xFLAG Δ*phoP790*::*kan*^+^) We amplified the Δ*rpoN730*::*kan*^+^ allele from strain MJG1763 with primers 5’ TAC AAG ACG AAC ACG TTA 3’ and 5’ TTT GGC AAA TTT GGC TGT 3’ and used recombineering (101) to insert this locus into strain MJG2191, generating strain MJG2200 (Δ*rpoN730*::*kan*^+^ *ptsN*-3xFLAG), which we then resolved (101) to generate strain MJG2202 (Δ*rpoN730 ptsN*-3xFLAG).

### Plasmid construction

Plasmid pRpoN was a gift from Dr. Joseph Wade (NY State Department of Health)(103). We amplified the *glnK* CDS (339 bp) plus 20 bp of upstream sequence from *E. coli* MG1655 genomic DNA with primers 5’ TTC GAA TTC ATT CTG ACC GGA GGG GAT CTA T 3’ and 5’ CTT AAG CTT TTA CAG CGC CGC TTC GTC 3’ and cloned it into the *Eco*RI and *Hin*dIII sites of plasmid pBAD18 (58) to generate plasmid pGLNK1. We amplified the *glnB* CDS (339 bp) plus 11 bp of upstream sequence and the *glmS* CDS (1830 bp) plus 25 bp of upstream sequence from *E. coli* MG1655 genomic DNA with primers 5’ TTT GGG CTA GCG AAT TCC AAG GAA TAG CAT GAA AAA GAT TGA 3’ and 5’ CAA AAC AGC CAA GCT TTT AAA TTG CCG CGT CGT C 3’ or 5’ TTT GGG CTA GCG AAT TCA CGA TAT AAA TCG GAA TCA AAA ACT ATG 3’ and 5’ CAA AAC AGC CAA GCT TTT ACT CAA CCG TAA CCG ATT TTG C 3’ and then inserted each gene between the *Eco*RI and *Hin*dIII sites of plasmid pBAD18 (amplified with primers 5’ AAG CTT GGC TGT TTT GGC 3’ and 5’ GAA TTC GCT AGC CCA AAA AAA C 3’) by *in vivo* assembly cloning (104) to generate plasmids pGLNB1 and pGLMS1, respectively. Plasmid pPPK33, encoding PPK with a C-terminal GAAEPEA peptide tag for affinity purification (105) between the *Nde*I and *Hin*dIII sites of plasmid pET-21b(+)(Novagen), was synthesized by GenScript.

We amplified the *gfpmut3* CDS (717 bp) (106) with primers 5’ TTG AAG GCT CTC AAG GAT GAG TAA AGG AGA AGA ACT TTT CAC TGG 3’ and 5’ CAG GGC AGG GTC GTT TTA TTT GTA TAG TTC ATC CAT GCC ATG TG 3’, the *aadA*^+^ gene and origin of replication of pCDFDuet-1 (Novagen) with primers 5’ AAC GAC CCT GCC CTG AAC C 3’ and 5’ CCT TGA GAG CCT TCA ACC CAG T 3’, and joined the resulting products by *in vivo* assembly (104), yielding plasmid pGFP3. We amplified the *mgtS* promoter (200 bp) from MG1655 genomic DNA with primers 5’ GGT TGA AGG CTC TCA AGG TGA TCA TTG CTG CGT GGG TGC TGA 3’ and 5’ AAG TTC TTC TCC TTT ACT CAT TAT TTG CCT CCG TTT ATG GTG TGG TGT G 3’, pGFP3 with primers 5’ ATG AGT AAA GGA GAA GAA CTT TTC ACT GGA G 3’ and 5’ CCT TGA GAG CCT TCA ACC CAG TC 3’, and joined the resulting products by *in vivo* assembly (104), yielding plasmid pGFP4.

We amplified the *ptsN* CDS (492 bp) plus 20 bp of upstream sequence from *E. coli* MG1655 genomic DNA with primers 5’ CTC TCT ACT GTT TCT CCA TAC CCG TTT TTT TGG GCT AGC GGC AGG TTC TTA GGT GAA ATT ATG ACA AAT AAT GAT ACA 3’ and 5’ TAT CAG GCT GAA AAT CTT CTC TCA TCC GCC AAA ACA GCC ACT ACG CTT CAT CCG GAG TAC CT 3’ and inserted it between the *Eco*RI and *Hin*dIII sites of plasmid pBAD18 by *in vivo* assembly cloning (104) to generate plasmid pPTSN1. We used single primer site-directed mutagenesis (107) to mutate pPTSN1 with primers 5’ CAA TGG TAT TGC CAT TCC GGA AGG CAA ACT GGA AGA AGA TAC 3’ or 5’ GGT ATT GCC ATT CCG GCG GGC AAA CTG GAA GAA G 3’. This yielded pPTSN2, containing a *ptsN*^C217G, T219A^ allele (encoding PtsN^H73E^), and pPTSN3, containing a *ptsN*^C217G, A218C, T219G^ allele (encoding PtsN^H73A^).

We amplified the dimerizing leucine zipper domain of GCN4 (105 bp) from *Saccharomyces cerevisiae* genomic DNA with primers 5’ TCC GGA TCC CTT GCA AAG AAT GAA ACA ACT TGA AG 3’ and 5’ ACC GGT ACC CGG CGT TCG CCA ACT AAT TTC T 3’ and cloned it into the *Bam*HI and *Kpn*I sites of plasmid pKT25 (73) to yield plasmid pGCN4zip1 and into the *Bam*HI and *Kpn*I sites of plasmid pUT18 (73) to yield plasmid pGCN4zip3. We amplified the *ppk* CDS with no stop codon (2084 bp) from *E. coli* MG1655 genomic DNA with primers 5’ CAG CTG CAG GGA TGG GTC AGG AAA AGC TAT ACA TCG 3’ and 5’ TCC GGA TCC TCT TCA GGT TGT TCG AGT GAT TTG 3’ and cloned it into the *Pst*I and *Bam*HI sites of plasmid pKNT25 (73) to yield plasmid pPPK12 or into the *Pst*I and *Bam*HI sites of plasmid pKT25 (73) to yield plasmid pPPK13. We amplified the *ptsN* CDS (493 bp) from plasmid pPTSN1 with primers 5’ CAC TGC AGG ATG ACA AAT AAT GAT ACA ACT CTA CAG CTT A 3’ and 5’ TGA ATT CGA CTA CGC TTC ATC CGG AGT AC 3’, amplified pUT18C (73) with primers 5’ GAA GCG TAG TCG AAT TCA TCG ATA TAA CTA AGT AAT ATG GTG 3’ and 5’ TAT TTG TCA TCC TGC AGT GGC GTT CCA C 3’, and joined the resulting products by *in vivo* assembly (104), yielding plasmid pPTSN5.

### *In vivo* polyphosphate assay

We extracted and quantified polyP from bacterial cultures as previously described (108). To induce polyP synthesis by nutrient limitation (16, 22), we grew *E. coli* strains in 10 ml rich medium (LB, LBQ, or LBGlcN) at 37°C with shaking (200 rpm) to A_600_=0.2-0.4, then harvested 1 ml samples by centrifugation, resuspended them in 250 µl of 4 M guanidine isothiocyanate, 50 mM Tris-HCl (pH 7), lysed by incubation for 10 min at 95°C, then immediately froze at -80°C. We also harvested 5 ml of each LB culture by centrifugation (5 min at 4,696 g at room temperature), rinsed once with 5 ml phosphate-buffered saline (PBS), then re-centrifuged and resuspended in 5 ml MOPS minimal medium (Teknova)(109) containing 0.1 mM K_2_HPO_4_, and 0.1 mM uracil and 4 g l^-1^ glucose (24). Where indicated, we replaced the glucose with 4 g l^-1^ arabinose, 4 g l^-1^ glucosamine, or 8 g l^-1^ glycerol. We incubated these cultures for 2 hours at 37°C with shaking, then collected additional samples as described above. For experiments involving arabinose-inducible plasmids, we added arabinose (2 g l^-1^) to both the rich and minimal media. We determined the protein concentrations of thawed samples by Bradford assay (Bio-Rad) of 5 µl aliquots, then mixed with 250 µl of 95% ethanol, applied to an EconoSpin silica spin column (Epoch Life Science), rinsed with 750 µl 5 mM Tris-HCl, pH 7.5, 50 mM NaCl, 5 mM EDTA, 50% ethanol, then eluted with 150 µl 50 mM Tris-HCl, pH 8. We brought the eluate to 20 mM Tris-HCl, pH 7.5, 5 mM MgCl_2_, 50 mM ammonium acetate with 1 µg of *Saccharomyces cereviseae* exopolyphosphatase PPX1 (110) in a final volume of 200 µl, incubated for 15 min at 37°C, then measured the resulting polyP-derived orthophosphate using a colorimetric assay (111) and normalized to total protein content. For all figures, we report polyP concentrations in terms of individual phosphate monomers.

### Quantitative RT-PCR

At the indicated time points after nutrient limitation, we harvested 1 ml of cells by centrifugation and resuspended in RNA*later* (ThermoFisher) for storage at -20°C. We extracted RNA using the RiboPure™ RNA Purification Kit for bacteria (Ambion) following the manufacturer’s instructions, including DNAse treatment to remove contaminating genomic DNA, then used the SuperScript^TM^ IV VILO^TM^ kit (ThermoFisher) to reverse transcribe cDNA from mRNA, following the manufacturer’s instructions and including a no-RT control for each reaction. We calculated changes in gene expression using the 2^-ΔΔCt^ method (112), using *yqfB*, whose expression does not change under these polyP induction conditions (22), as an internal expression control.

### PPK overexpression and purification

C-tagged PPK was overexpressed and purified by a modification of a previously published protocol (17). 50-ml overnight cultures of BL21(DE3) containing pPPK33 were subcultured into 1 l of Protein Expression Media (PEM; 12 g l^-1^ tryptone, 24 g l^-1^ yeast extract, 4% v/v glycerol, 2.314 g l^-1^ KH_2_PO_4_, 12.54 g l^-1^ K_2_HPO_4_) supplemented with 10 mM MgCl_2_ and 100 µg ml^-1^ ampicillin. The culture was grown at 37°C with shaking until the A_600_ = 0.8, then shifted to 20°C and cooled for 1 hour. Following the cool-down period, PPK expression was induced by the addition of 150 µM isopropyl *β*-D-1-thiogalactopyranoside (IPTG). Overexpression was allowed to proceed overnight at 20°C with shaking. The overexpression culture was pelleted at 6000 g, resuspended in 100 ml of Buffer A (50 mM Tris-HCl pH 7.5, 10% w/v sucrose) with 300 µg ml^-1^ lysozyme, and incubated on ice for 45 min. The digest was pelleted at 16,000 x g for 10 minutes, then the pellet was resuspended in 50 ml of Buffer B (Buffer A + 5 mM MgCl_2_ + 30 U ml^-1^ Pierce Universal Nuclease + 1 Pierce Protease Inhibitor cocktail tablet) and lysed by sonication (5 s on, 5 s off for 5 min at 50% amplitude). The sonicated lysate was pelleted at 20,000 g for 1 hour at 4°C, and the pellet was resuspended in 25 ml of C-tag Binding Buffer (20 mM Tris-HCl pH 7.4) plus solid KCl to 1 M final concentration. 1 M Na_2_CO_3_ was added at a 1:10 dilution to the resuspension, and the salt extraction was incubated at 4°C for 30 min with stirring. Following incubation, the solution was sonicated in 5 s pulses for 2 min, then pelleted at 20,000 g for 1 hour at 4°C. The supernatant was diluted 1:1 with cold H_2_O and loaded onto a C-tag Affinity Column (ThermoFisher) equilibrated with C-tag Binding Buffer. The column was washed with 10 column volumes of C-tag Binding Buffer and PPK was eluted with a gradient of 0-100% C-tag Elution Buffer (20 mM Tris-HCl pH 7.4, 2 M MgCl_2_) with an ÄKTA^TM^ Start FPLC (Cytiva Life Sciences). Fractions containing pure PPK were pooled and dialyzed against PPK Storage Buffer (20 mM HEPES-KOH pH 8.0, 150 mM NaCl, 15% glycerol, 1 mM EDTA) at 4°C and stored at -80°C.

### *In vitro* assay of PPK activity

We determined the specific activity for polyP synthesis by PPK as previously described (23). Reactions (125 µl total volume) contained 5 nM PPK, 50 mM HEPES-KOH (pH 7.5), 50 mM ammonium sulfate, 5 mM MgCl_2_, 20 mM creatine phosphate, and 60 µg ml^- 1^ creatine kinase. Where indicated, reactions also contained 100 mM *L*-glutamate (pH 7.5), 10 mM *L*-glutamine (pH 7.5), 1 mM fructose-6-phosphate, 1 mM glucosamine-6-phosphate, or 5 or 10 mM *α*-ketoglutarate (pH 7.5), concentrations chosen to represent the high end of the physiological range for each compound in *E. coli* (51). We prewarmed reactions to 37°C, then started them by addition of MgATP to a final concentration of 6 mM. We removed aliquots (20 µl) at 1, 2, 3, and 4 minutes and diluted them into 80 µl of a stop solution containing 62.5 mM EDTA and 50 µM DAPI (4’,6-diamidino-2-phenylindole) in black 96-well plates, then measured steady-state polyP-DAPI fluorescence of these samples (ex. 415 nm, em. 600 nm) (15) in an Infinite M1000 Pro microplate reader (Tecan Group, Ltd.). We determined the polyP content of each sample (calculated in terms of individual phosphate monomers) by comparison to a standard curve of commercially available polyP (Acros Organics) (0 - 150 µM) prepared in the buffer described above containing 6 mM MgATP and calculated rates of polyP synthesis by linear regression (Prism 9, GraphPad Software, Inc.).

### Bacterial two-hybrid protein interaction assay

We assessed protein interactions *in vivo* using the BACTH procedure (73). Briefly, we grew derivatives of *E. coli cya* strain BTH101 containing plasmids expressing fusions of proteins of interest to the T18 or T25 complementary fragments of *Bordetella pertussis* adenylate cyclase overnight in LB and then evaluated *β*-galactosidase activity of these strains either by spotting overnight cultures on LB plates containing ampicillin, kanamycin, 0.5 mM IPTG, and 40 µg ml^-1^ 5-bromo-4-chloro-3-indolyl-*β*-D-galactopyranoside (X-Gal) and incubating for 2 days at 30°C or, for quantitative measurements, using a single-step assay (113): after 24 h of growth at 37°C in LB broth containing ampicillin, kanamycin, and 0.5 mM IPTG we harvested 80 µL of cells by centrifugation, resuspended them in 200 µL of 60 mM Na_2_HPO_4_, 40 mM NaH_2_PO_4_, 10 mM KCl, 1 mM MgSO_4_, 36 mM *β*-mercaptoethanol, 1.1 mg ml^-1^ *ortho*-nitrophenyl-β-galactoside (ONPG), 1.25 mg ml^-1^ lysozyme and 6.7% PopCulture reagent (Novagen) in a 96-well plate, and then measured A_600_ and A_420_ over time at 24°C in a Tecan M1000 Infinite plate reader. We calculated Miller Units according to the formula (1000 x (A_420_ / min)) / (initial A_600_ x culture volume (ml)).

### Quantitative Western blotting

*E. coli* strains with chromosomal *ptsN*-3xFLAG fusions were grown and stressed by nutrient limitation as described above. At the indicated time points, 1-ml aliquots were harvested by centrifugation and resuspended in 100 µl of 50 mM Tris-HCl (pH 8), 150 mM NaCl, 1% Triton X-100 containing 1X HALT protease inhibitor cocktail (ThermoFisher), then incubated at 95°C for 10 min to lyse the cells. Lysates were stored at -80°C until use. Aliquots of each sample were thawed on ice, mixed 1:1 with fresh lysis buffer, then mixed 4:1 with reducing loading dye (250 mM Tris-HCl pH 6.8, 10% SDS, 0.008% bromophenol blue, 40% glycerol, 2.8 M *β*-mercaptoethanol). Western blots were prepared and analyzed as described previously (114, 115) with few exceptions. Briefly, lysate samples were loaded on an AnykDa Stain-Free SDS-PAGE gel (BioRad Cat #4568126) and run until the dye front neared the bottom of the gel. Gels were then transferred to a PVDF membrane (BioRad Cat # 1620174) using a TurboBlot semi-dry transfer system, then blocked in StartingBlock T20 TBS blocking buffer (Thermo Cat #37543) overnight. Blots were blocked for 30 min at room temperature, then incubated in a 1:25,000 dilution of rabbit anti-RecA antibody (Abcam Cat #ab63797) in the blocking buffer for one hour at room temperature. Blots were washed in three times in TBST, then incubated 1 hour at room temperature in blocking buffer containing a 1:10,000 dilution of goat anti-Rabbit IgG H+L HRP-conjugated (Abcam Cat #ab63797). Blots were washed again in 3X TBS-T then incubated in blocking buffer containing a 1:5000 dilution of rabbit Anti-DDDDK HRP conjugated antibody (Abcam Cat #ab2493). Blots were washed three times with TBST and once with TBS, then developed using the BioRad Clarity ECL Substrate kit (Cat # 1705061) and imaged on a BioRad Gel Doc. Images were analyzed in ImageJ (116) by taking same-area measurements of each RecA band or PtsN band, blank correcting the mean gray values (signal) using same-area measurements matched to the respective band, and normalizing the PtsN signal to RecA signal.

### Statistical analyses

We used GraphPad Prism version 9.2 (GraphPad Software) to perform statistical analyses, including two-way repeated measures ANOVA with Holm-Sidak’s multiple comparison tests. Repeated measures ANOVA cannot handle missing values, so we analyzed data sets with samples having different *n* numbers (*e.g.* Fig. 1C) with an equivalent mixed model which uses a compound symmetry covariance matrix and is fit using Restricted Maximum Likelihood (REML) (without Geisser-Greenhouse correction).

### Data availability

All strains and plasmids generated in the course of this work are available from the authors upon request.

## ACKNOWLEDGEMENTS

This project was supported by NIH grant R35GM124590.

